# Nest aggregations of wild bees and apoid wasps in urban pavements: a “street life” to be promoted in urban planning

**DOI:** 10.1101/2021.12.15.472743

**Authors:** Grégoire Noël, Violette Van Keymeulen, Yvan Barbier, Sylvie Smets, Olivier Van Damme, Gilles Colinet, Julien Ruelle, Frédéric Francis

**Affiliations:** Functional and Evolutionary Entomology, Gembloux Agro-Bio Tech, University of Liège, Passage des Déportés 2, 5030 Gembloux, Belgium; Service public de Wallonie, Département du Milieu Naturel et Agricole, Avenue de la Faculté, 22, B-5030 Gembloux, Belgium; Belgian Road Research Centre (BRRC), Boulevard de la Woluwe, 42, B-1200 Woluwé-Saint-Lambert, Belgium; Soil-Water-Plant Exchanges, Gembloux Agro-Bio Tech, University of Liège, Passage des Déportés 2, 5030 Gembloux, Belgium; Département Développement Nature et Agriculture – Bruxelles Environnement (BE), Site de Tour & Taxis, Avenue du Port 86C / 3000, 1000 Bruxelles, Belgium

**Keywords:** Anthophila, Apoidea, nesting behaviour, urban ecology, urban ecosystem, urban conservation

## Abstract

In the last 10 years, knowledges of wild bees and apoid wasps’ community dynamics have gained interest in urban ecology focusing on the availability of floral resources in cities. Although understudied, the urban environment impacts the conditions of their nesting sites. Recent observations in the Brussels-Capital Region (Belgium) showed that urban pavements can be a novel nesting opportunity for Hymenoptera ground-nesting species such as wild bees and apoid wasps. Here, using citizen science, we investigated the richness of ground-nesting species living under urban pavements, the preferences of the sidewalk joint size related to ground-nesting species size and for sidewalk type or for soils texture under the pavements on the nesting site selection. A total of 22 species belonging to 10 Hymenoptera families of wild bees and digger wasps with their associated kleptoparasites were identified on 89 sites in Brussels. Sandstone setts or concrete slabs with an unbound joint size around 1 cm were found to be best suitable urban pavements for the ground-nesting species. The soil texture under the pavement was highly sandy among our samples. Finally, we also suggest engineering management guidelines to support bee and wasp species nesting under urban pavement in highly urbanized areas. Such observations pave the way for much research in the field of urban ecology to conceive multifunctional pavement promoting biodiversity.

## Introduction

In 2007, the United Nations estimated that the proportion of human population in cities had overtaken rural ones. Since then, the process of urban migration accelerated and more than 67% of humanity is expected to concentrate in cities by 2050 (United Nations 2019). Urban and suburban areas are often perceived as being poor sites for biodiversity. Indeed, urban ecosystems cause many disturbances that can drastically influence the evolutionary and ecological dynamics of populations (Alberti 2015; Alberti et al. 2017). Cities not only alter biodiversity by reducing the number of native species but also act as a selective agent and determine which species can live while inducing eco-evolutionary changes in organisms (Shochat et al. 2010). For example, many organisms, including birds, fish, mammals, plants and arthropods, adapt to urban environments by changing their physiology, morphology and behaviour (Hendry et al. 2008; Palkovacs and Hendry 2010; Johnson et al. 2018).

Hymenopteran communities also seem to be sensitive to landscape urban conversion (Buczkowski and Richmond 2012; Geslin et al. 2016; Corcos et al. 2019; Theodorou et al. 2020a). The amount of floral resources becomes scarcer under the pressure of urban fragmentation but also with soil imperviousness due to increasing pavement that makes nesting sites inaccessible for ground-nesting Hymenopteran species (Burkman and Gardiner 2014; Harrison and Winfree 2015; Geslin et al. 2016; Wenzel et al. 2020; Ayers and Rehan 2021). Particularly, some research has shown a decrease in the richness of urban bee communities as well as a decrease in their size along an increasing urbanization gradient (Ahrné et al. 2009; Fortel et al. 2014; Eggenberger et al. 2019). Paradoxically, recent studies have shown that cities can also serve as refuges for wild bee communities (Baldock et al. 2015; Hall et al. 2017; Theodorou et al. 2020b). Several explanations have been put forward for this: (i) the rate and coverage of biocidal particles in cities is usually lower than in the surrounding countryside, (ii) the urban heat island phenomenon favours the persistence of thermophilic species, (iii) the heterogeneity of urban patches allows the coexistence of a large diversity of habitats and then a wide variety of associated ecological niches, and (iv) urban parks, gardens and other green spaces generally allow for sufficient floral resources distributed throughout the year (Fortel et al. 2016; Wenzel et al. 2020; Fenoglio et al. 2021). More than half of the 403 species of Belgian wild bees have a ground-nesting behaviour (Drossart et al. 2019). These solitary bees build below ground nest usually consisting of a main gallery branched into secondary galleries containing the larval cells in which they store food resources (mixture of pollen and nectar) before laying eggs (Malyshev 1935; Michener 2007). These species belong to Andrenidae, Melittidae as well as a majority to Halictidae and Colletidae families (Danforth et al. 2019; Drossart et al. 2019). Currently, the state of knowledge about adaptations of these ground-nesting bees in urban environments remains patchy especially related to their nesting strategies (Wenzel et al. 2020; Antoine and Forrest 2020; Ayers and Rehan 2021).

In urban environment, apoid wasps - including recently up-ranked families (e.g., Philantidae, Psenidae, Bembicidae, Pemphredonidae) by Sann et al., (2018) - play also important ecological roles : pollination and predatory biocontrol. Adults behave as flower-visitors and capture insect or spider prey as predator to feed their offspring (Bitsch and Leclercq 1993). In Belgium, 199 species of apoid wasps were recorded and more than half nests under the ground (i.e., 107 spp.) (Pauly 1999; Rasmont and Haubruge 2002). Furthermore, some taxa can show collective nesting strategy in nest aggregation such as some *Cerceris* spp. (Willmer 1985; Polidori et al. 2006). The known deterioration of their populations is mainly due to changes in habitat losses that impacts favourable ground nesting sites and sufficient food resources (Gauld et al. 1990; Day 1991; Zanette et al. 2005; Christie and Hochuli 2009). Moreover, their ability to locate preys in large landscape is disrupted by the habitat fragmentation caused by urban matrix (Kareiva 1987) leading to change in the community structure of apoid wasps in cities (Christie and Hochuli 2009; Burkman and Gardiner 2014). However, to our knowledge, there is still no study mentioning nesting adaptation regarding to urban matrix for apoid wasps.

Urbanization leads to a considerable increase in impervious surfaces (buildings, sidewalks, roads…), which limit nesting opportunities on natural substrates (Cane 2005) and would lead to competition for these sites among ground-nesting bees and apoid wasps. Nevertheless, recent observations in the Brussels-Capital Region (BCR, Belgium) showed that urban pavements have become a novel nesting opportunity for certain ground-nesting species (Pauly 2019a). A sandy mound indicates the presence of these ground-nesting species (Fig. 1a). Indeed, the drainage of flexible pavements, the recurrent use of sandy substrate under the paving materials and the thermal capacity of the pavements could probably contribute to the potential of pavements to host ground-nesting bees and wasps, providing them an interesting shelter in an urban environment. The joint size could have a direct impact on the nesting ability of the bees and wasps: when joint size is lower than their thorax size, it does not allow digging a gallery. Based on the scientific literature on variables that influence nest site selection in natural environments (Stephen 1960, 1965; Osgood Jr 1972; Cane 1991, 2015; Potts and Willmer 1997; Wuellner 1999; Sardiñas and Kremen 2014; Anderson and Harmon-threatt 2016; Harmon-Threatt 2020; Nichols et al. 2020), we hypothesized the following variables that influenced nest site selection in pavements for urban environments: the nature of the jointing material, the type of pavement and the location of the nest on the pavement. As an entry zone to the subsurface material, the nature of the jointing material defines the hardness of the substrate and would therefore influence the ability of bees to tunnel into it, acting as a filter in nest site selection.

**Fig. 1.**
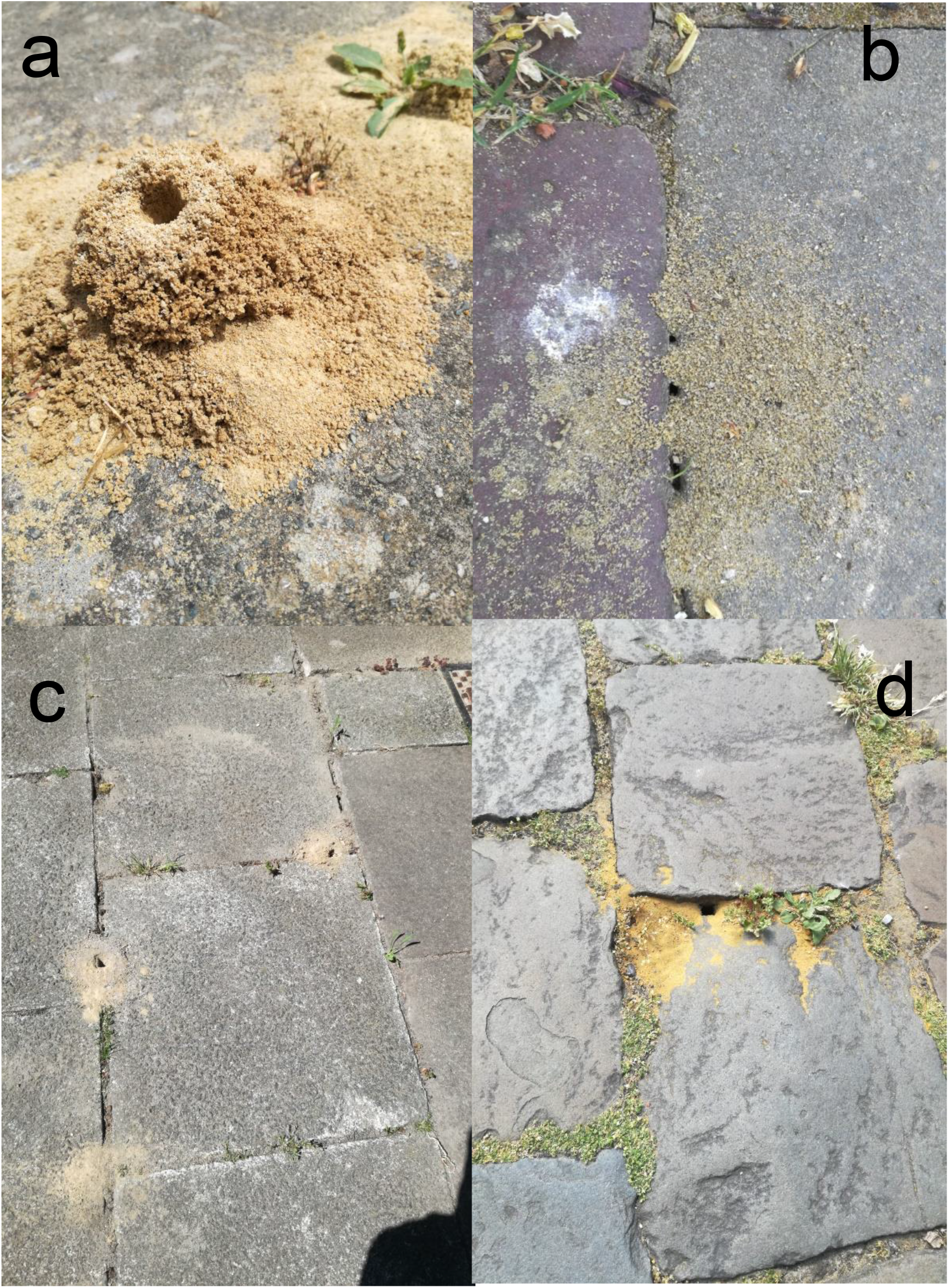
Nest pictures on pavements. Sandy mound (Auderghem, Brussels) (**a**). Ant nest (Anderlecht, Brussels) (**b**). Degraded rigid joints of concrete slabs (Schaerbeek, Brussels) (**c**). Unbound joints of sandstone setts (Schaerbeek, Brussels) (**d**). Pictures by Grégoire Noël.

As a pioneering study in urban ecology of Hymenoptera ground-nesting species, we addressed the following questions regarding wild bee and apoid wasp community according to their nesting preferences up and under the pavements: (i) what is the ground-nesting community living under the BCR pavements ? (ii) Is there a link between ground nesting species size and joint size ? (iii) Does the pavements type have an impact on it ? (iv) How is the soil texture under pavement selected by ground-nesting species ? In an attempt to answer these questions, we developed a citizen science pipeline to identify their nesting sites followed by an appropriate methodology to characterize the selected nesting sites. After species identification, an analysis of associated edaphic preferences was performed for pavements selection. Finally, the implications of our results were discussed regarding to the new challenges in urban pavement design that should promote ground-nesting insects.

## Material and methods

### Identification of the potential study sites

Only 4 nesting sites on the pavements had been identified and confirmed in BCR by Pauly in 2019a. To increase our sampling effort, we developed a crowdsourcing method based on citizen science (Newman et al. 2017). Then, we created and actively disseminated a participatory survey to BCR citizens on social networks in collaboration with the communication department of Brussels Environment and other key actors in BCR conservation (e.g., regional and local institutions …). The coding form was written on February 10, 2020 and put online on March 11, 2020 before the first emergence of the potential hymenopteran insects (Pauly 2019a). In order to ensure and increase the quality of the observations received through the form (Aceves-Bueno et al. 2017), several elements were set : (i) the “*commune*” field only allows the encoding of BCR municipality in order to avoid encoding data outside our study field, (ii) the street name and the building number are two separate mandatory fields in order to ensure accurate locations and to reduce the search field for subsequent field validations, (iii) the date of the last observation of the nest, (iv) the potential picture of the nest or the insect in presence to pre-validate that the observation took place on a pavement and is related to our taxa of interest.

### Validation of the study sites

The dataset curation from the coding form was started in mid-March 2020 and continued throughout the participatory survey until mid-July 2020. We removed observations unrelated to taxa and location of our study sampling strategy (e.g., ants, cavity-nesting bees, outside of BCR …). We validated study sites on the field from April 05, 2020 to July 20, 2020 during highly active flying periods which refers to sunny days with clear sky, little wind (less than 15 km/h) and minimum daily temperature of 15°C between 09:00 AM and 05:00 PM (Ahrné et al. 2009; Fortel et al. 2014). Once on site, we examined the pavement cover for 30-45 minutes. The site was validated if: (i) a bee or wasp showed evidence of entry or exit activity in its nest between the slabs and/or (ii) a cuckoo species (i.e., cuckoo bee or wasp) patrolled near the sandy mound on the pavement. The ant nests (Hymenoptera: Formicidae) were recognizable by several clues: the sand grains were spread over the surface of the slab and did not form a mound, the nest had several visible contiguous entrances, and the introduction of a twig into one of the entrances revealed ant activity attempting to protect their territory (Fig. 1b).

### Data collection on study sites

The number of nests was estimated through the counting of sandy mounds. Within each of the selected sites, each morphologically distinct specimen was captured and killed *in situ* with the introduction of a drop of ethyl acetate for later species identification. In some cases, specimens were stored in a freezer at -20°C directly after field survey.

The joint size was measured on 6 nests randomly chosen within a site using a millimetre lath perpendicular to the joints and passing through the nest entrance. When more than one ground-nesting species was on site, we took measurements randomly on site, without distinction between species. The joint structure variable was implemented in the database as a nominal qualitative variable with 2 modalities: degraded rigid joint (Fig. 1c) / unbound joint (Fig. 1d). The type of pavement was evaluated on the basis of photographs and classified according to 3 modalities: concrete slabs (Fig. 1c) / sandstone setts (Fig. 1d) / other including ceramic slabs, concrete pavement, limestone (Belgian blue stone) and porphyry setts. Finally, the position of the nest on the pavement was implemented as a qualitative variable according to the following nomenclature: pavement / front of house / internal yard / road with car traffic / other.

Sand of the mounds was collected from random 3-10 nest entrances (max. 50g) on pavements as corresponding to the soil layer excavated by insects digging activity under pavements (Fig. 1a). Regarding to preliminary results of the substrate texture under pavements, sandy mounds seem to be a good proxy of soil texture (see in supplementary information S1).

### Laboratory data collection

All collected specimens were prepared for identification using Mouret *et al*., (2007) protocol. We used several identification keys to define bee and wasp species (Bitsch and Leclercq 1993; Bitsch et al. 1997, 2007; Falk 2015; Pauly 2019b). All identified bee specimens were checked via the reference collections of the Functional and Evolutionary Entomology Laboratory (ULiège), notably the collections of Alain Pauly for captured Halictidae species and the collections of Jean Leclercq for apoid wasp species. All apoid wasp species as well as chrysid species were confirmed or identified by Yvan Barbier (DEMNA).

Each of the sandy mound samples were weighed using a precision balance and passed through a sieve shaker (Haver & Boecker VWR brand) for 10 minutes at an oscillation amplitude of 1 mm through five sieves with mesh sizes of 1 mm, 500 μm, 200 μm, 100 μm and 50 μm. The choice of sieves was considered to allow for the distinction of sands from clays and silts (50 μm threshold) and to differentiate between very fine, fine, medium and coarse sands. The particles retained by each sieve were then weighed and their value converted to a percentage of the total sample volume. This conversion eliminates weight variations due to sample moisture and creates a common basis for comparison between samples of varying weights. Although the particle rate of silts and clays was not distinguished, we can approximate the average and extreme textures of the collected mounds through the textural triangle by also halving the remaining percentages between the silt or clay classes. If more than one species (bee or wasp) was present at a site, we labelled the sample by the dominant ground-nesting species based on our filed observations.

### Mapping and statistical analysis

All analysis were performed on R software environment (R Core Team 2020). The validated sites were mapped using *mapview* R package (Appelhans et al. 2019). For the joint size, we used an ANOVA after descriptive statistical analysis of the data to compare joint size measurements among selected ground-nesting species and among their respective families. For statistical analysis, we removed cuckoo species given their nesting strategy and the specimens from Bembicidae, Crabronidae, Psenidae and Pemphredonidae families by lacking of data (Table 1). A *post-hoc* Tukey test with adjustment of multiple comparisons was applied to compare mean pair of joint size. We also measured the species size as well as the inter-tegular distance (ITD) - distance between the two wing insertions - of only female individuals which are a proxy for their size (Kendall et al. 2019) using a digital caliper (Electronic Digital Caliper). The average mean joint size per site was therefore attributed to the corresponding species. We evenly distributed the ITD and joint size measurement by species present on site. A linear regression was performed to test the relation between species size and average mean joint size.

**Table 1.**
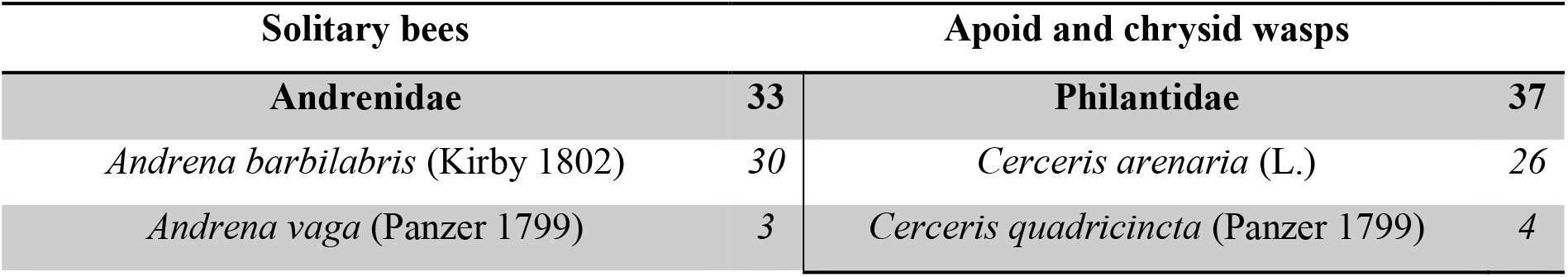
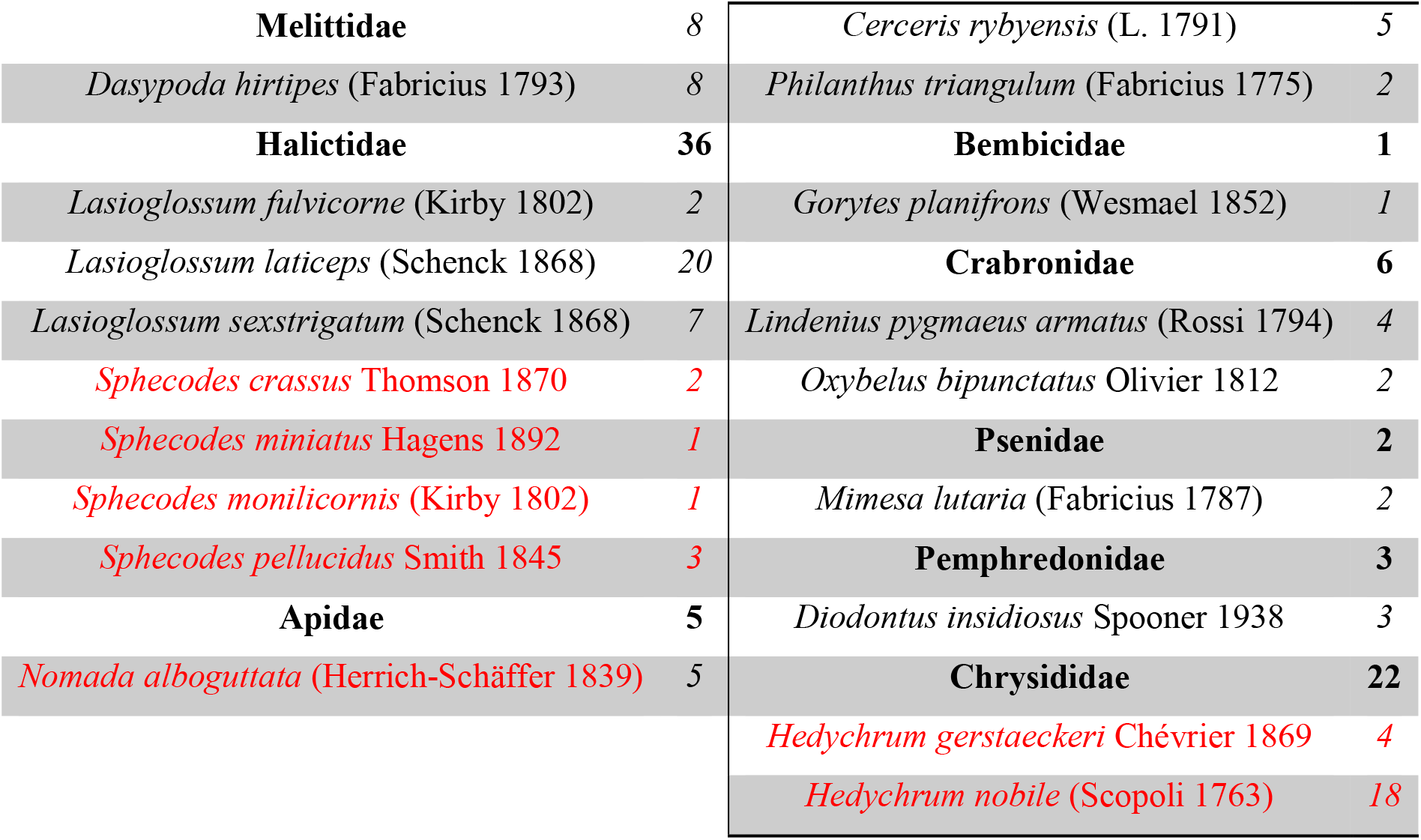
List of apoid and chrysid families and species collected at all sampling sites. The specific abundance is presented on the right side of each species. Species highlighted in black and red corresponded to ground-nesting and to bee - chrysid cuckoo species respectively. The apoid wasp families were defined according to Sann et al., (2018).

In relation to the particle size analysis, a principal component analysis (PCA) was performed to detect whether there were similarities in particle size preferences the ground-nesting species and between their respective families (as shown in Table 1) using *factoextra* (Kassambara and Fabian 2020) and *FactoMineR* (Lê et al. 2008) R packages. The graphs were drawn using *ggplot2* R package (Wickham 2016).

## Results

### Participatory survey

From March 11 to July 20, 2020, 163 observations were recorded on the survey form for BCR. The response frequency to the form varied around an average of 1.00 ± 2.00 (mean ± SD) entry per day. The municipalities of Woluwé-Saint-Lambert, Watermael-Boitsfort and Schaerbeek form the pack head, while Molenbeek-Saint-Jean, Jette and Ganshoren completed it (Fig. S1). There was no encoding from citizens of Saint-Josse ten-Noode and Koekelberg. Between April 5 and July 31, 2020, we characterized 89 sites throughout BCR that fullfilled the validation criteria (Fig. 2). Ixelles, Watermael-Boitsfort and Uccle municipalities were the most sampled (Fig. S2).

**Fig. 2.**
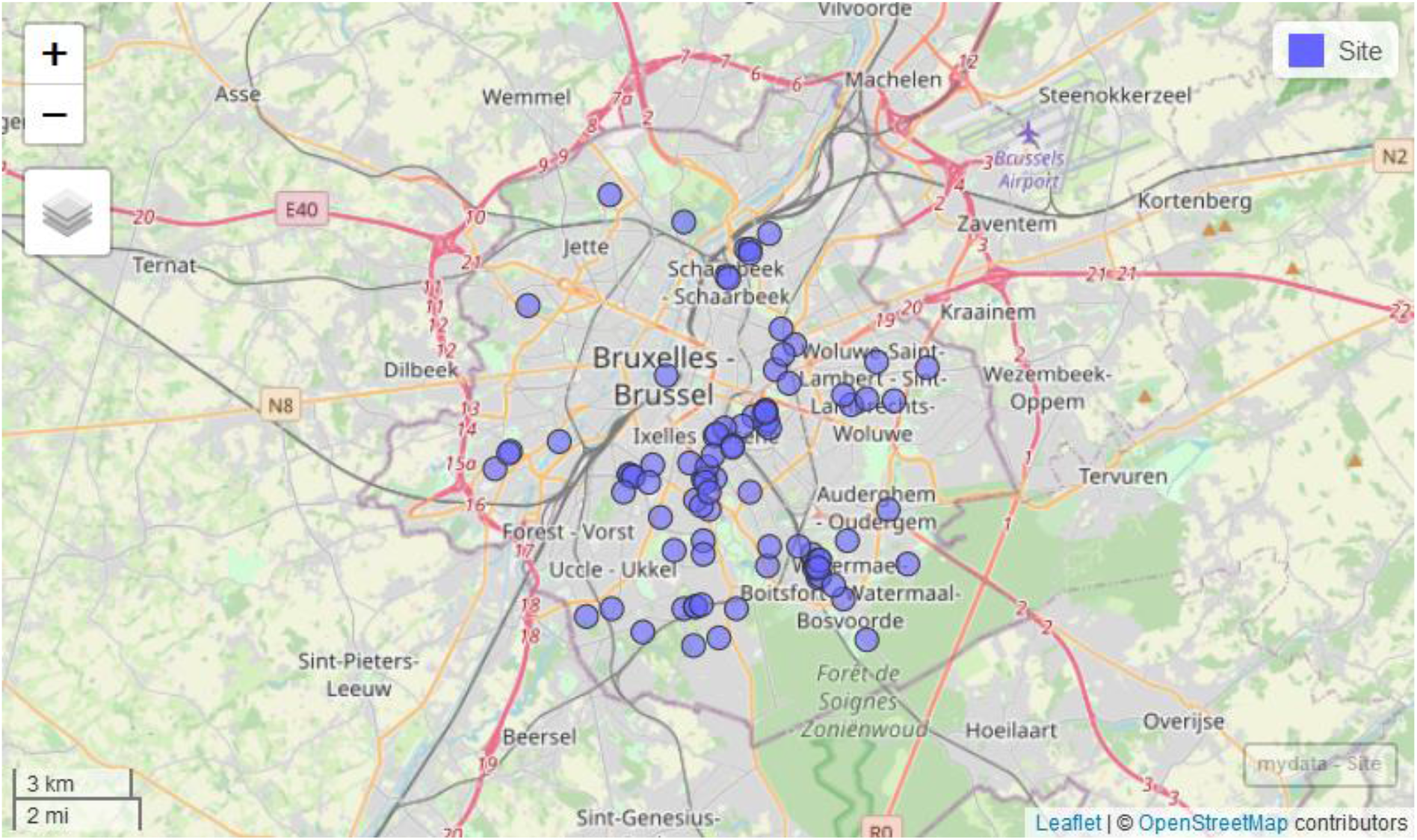
Distribution map of the validated study sites in Brussels Capital Region (N=89)

### Species recorded

We collected 153 specimens belonging to 22 species including 11 solitary bee species, 9 apoid wasp species and 2 chrysid species (Table 1).

The most abundant species found on studied sites were *A. barbilabris, C. arenaria, L. laticeps* and *H. nobile*. In addition, we also collected the cleptoparasite and parasitoid species, namely *N. alboguttata, Sphecodes* spp., *H. gerstaeckeri*, and *H. nobile* (Table 1). Most sites had a single ground-nesting species (excluding cleptoparasite and parasitoid species). However, some sampled sites showed that a co-occurrence between several nesting species is possible between solitary wasps and solitary bees but also between different bee species and between different wasp species (Fig. S3).

### Joint Size Analysis

After dataset curation, a total of 398 joint measurements were made at 69 validated sites on only 10 ground-nesting species (Fig. 3). The average joint size for all species was 1.08 cm ± 0.57 cm with a maximum measured at 3.00 cm and a minimum at 0.20 cm. Details by species were given in Table S1. A significant difference among the mean joint size between different ground-nesting species was observed (df = 9; F = 1.97; P = 0.041). However, after adjustment of multiple comparisons, no pairs of species differing in their joint size were detected (P > 0.05). After exclusion of some sites due to degraded individuals (without ITD measurements), the linear regression was performed on the average joint size and ITD size on 80 observations of ground-nesting specimens. ITD cannot explain the selection of joint size among the species (F = 0.16; df = 78; P = 0.69; Fig. 4).

**Fig. 3.**
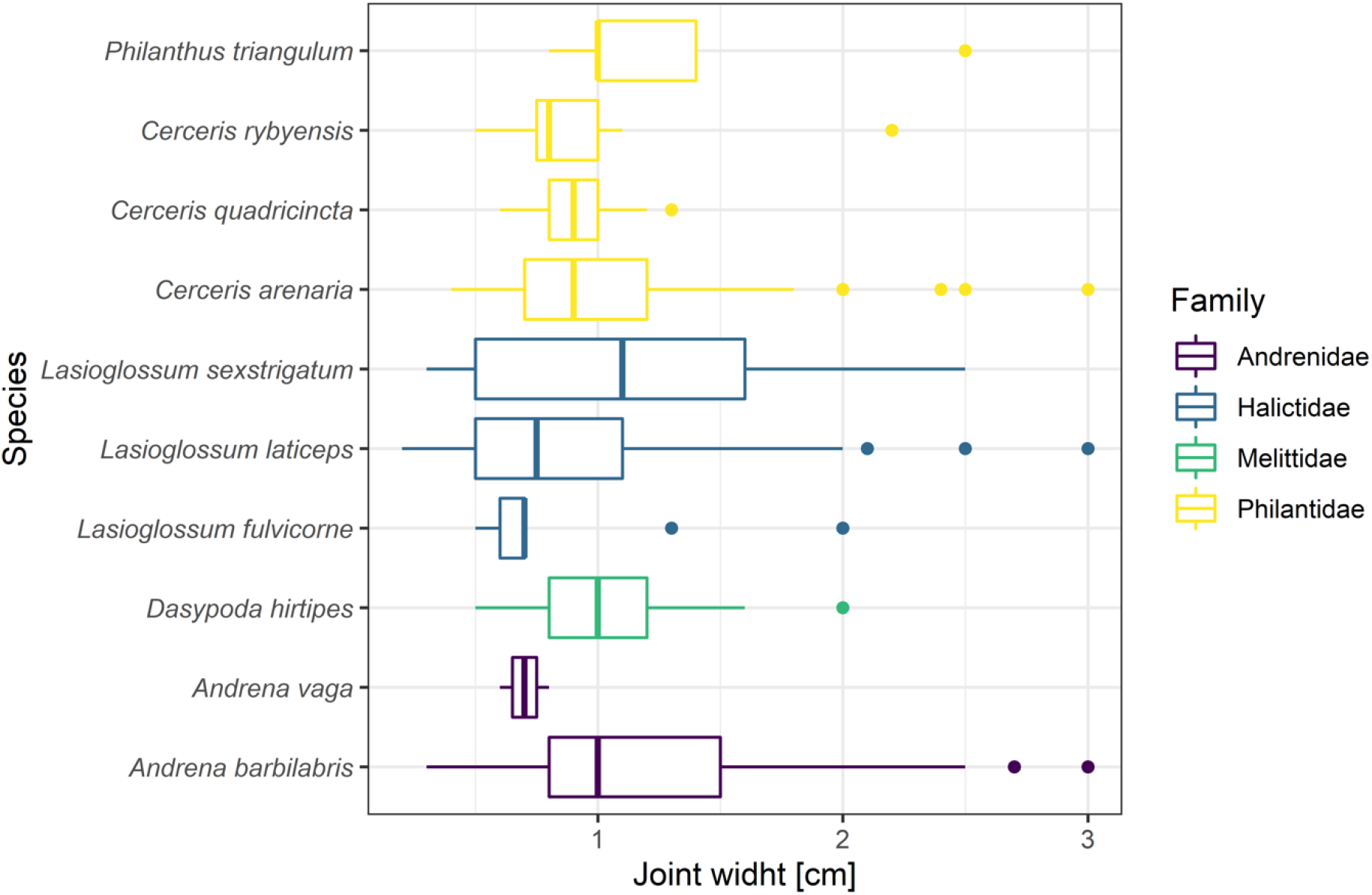
Distribution of joint sizes for nest entrances (in cm) according to different ground-nesting species and their respective families.

**Fig. 4.**
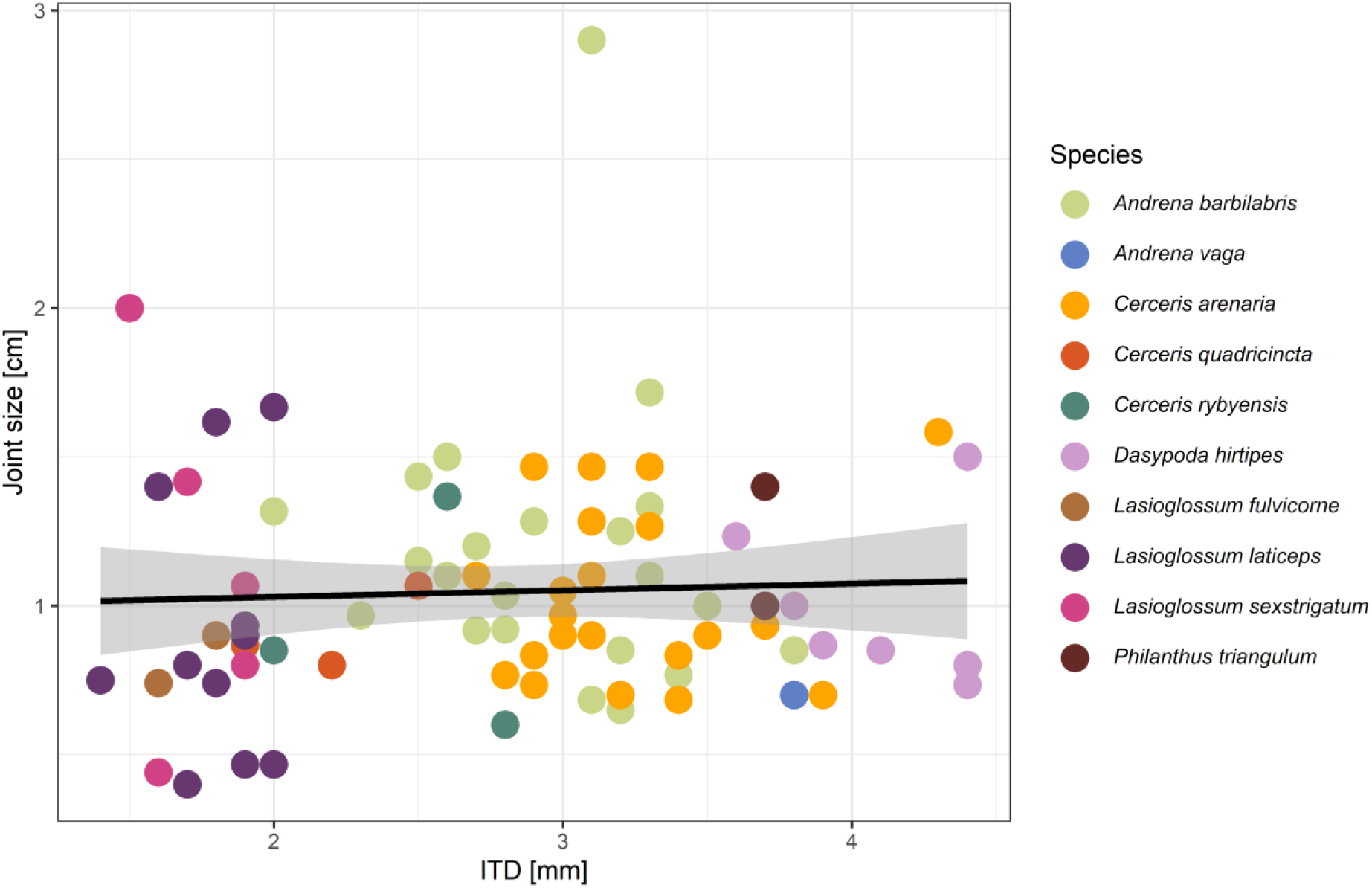
Linear regression of inter-tegular distance average (ITD in mm) and joint size average (in cm). Colors corresponded to different ground-nesting species. Grey shade area indicates 95% confidence interval region computed from means.

### Joint structure and pavement type

As some data were not taken into account due to misfit with our classification (e.g., hybrid pavement …), a total of 79 sites were characterized. The joints were mostly unbound and account for 80% of the sites encountered whereas 20% of the joints were characterized as rigid and degraded, leaving openings for ground-nesting species to dig and nest (Fig. 5a). As for the composition, pavements were mainly composed of concrete slabs (40 sites) and sandstone setts (29 sites) (Fig. 5b). The other sites were respectively composed of sandstone or limestone paving stones (3 sites), concrete paving blocs (4 sites), ceramic paving flags (1 site), porphyry setts (1 site) and blue stone elements (1 site). Regarding to the location of nests on pavements (Fig. 5c), most of them were located on sidewalks (53 sites) while some were located on roads (3 sites) and in the internal courtyards of houses (8 sites). Some sites were located at the level of house steps only (11 sites) or spilled sometimes over the sidewalks (2 sites). Only one site was characterized on stair steps and another one was characterized with an overflow of the ground-nesting aggregation from the embankment to the sidewalk.

**Fig. 5.**
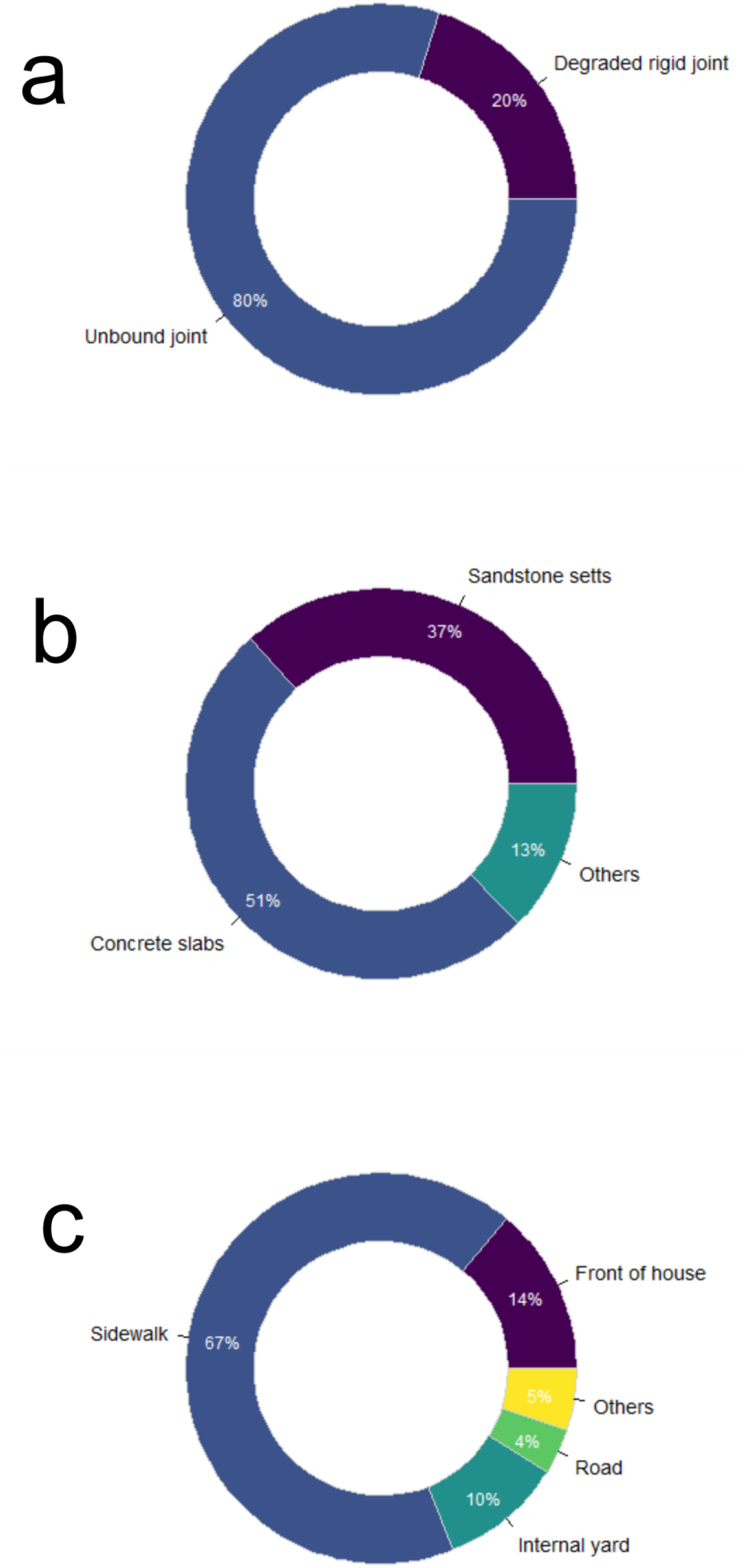
Distribution (%) from 79 study sites of joint types (**a**), kinds of urban pavement on which nests were located (**b**) and nest locations on urban pavements by site (**c**).

### Soil texture analysis

A total of 53 sandy mounds samples, which reached enough soil matter *in situ*, were analysed for grain size. The particle size analysis method did not allow us to distinguish between clay and silt composition. The sandy fraction of the samples was always higher than 85% and the silt and clay fractions were always lower than 10%, which placed all the samples from mounds as sandy and homogeneous texture (Fig. S4).

The samples were on average composed of 2.91% of particles larger than 1mm in diameter (i.e., very coarse sands and 2.61% smaller than 50 μm (i.e., clays and silts). Particles with a diameter of 500 μm - 200 μm (medium sands) were the most represented in the samples with a proportion of 41.71%. The samples contained on average 7.47% of particles of class 1 mm – 500 μm (coarse sands), 13.40% of particles with a diameter of 100 μm – 50 μm (very fine sands) and 31.90% of 200 μm – 100 μm (fine sands) (Fig. S5). Our sandy samples did not allow us to distinguish groupings or discontinuities according to the ground-nesting species or their respective families (Fig. 6).

**Fig. 6.**
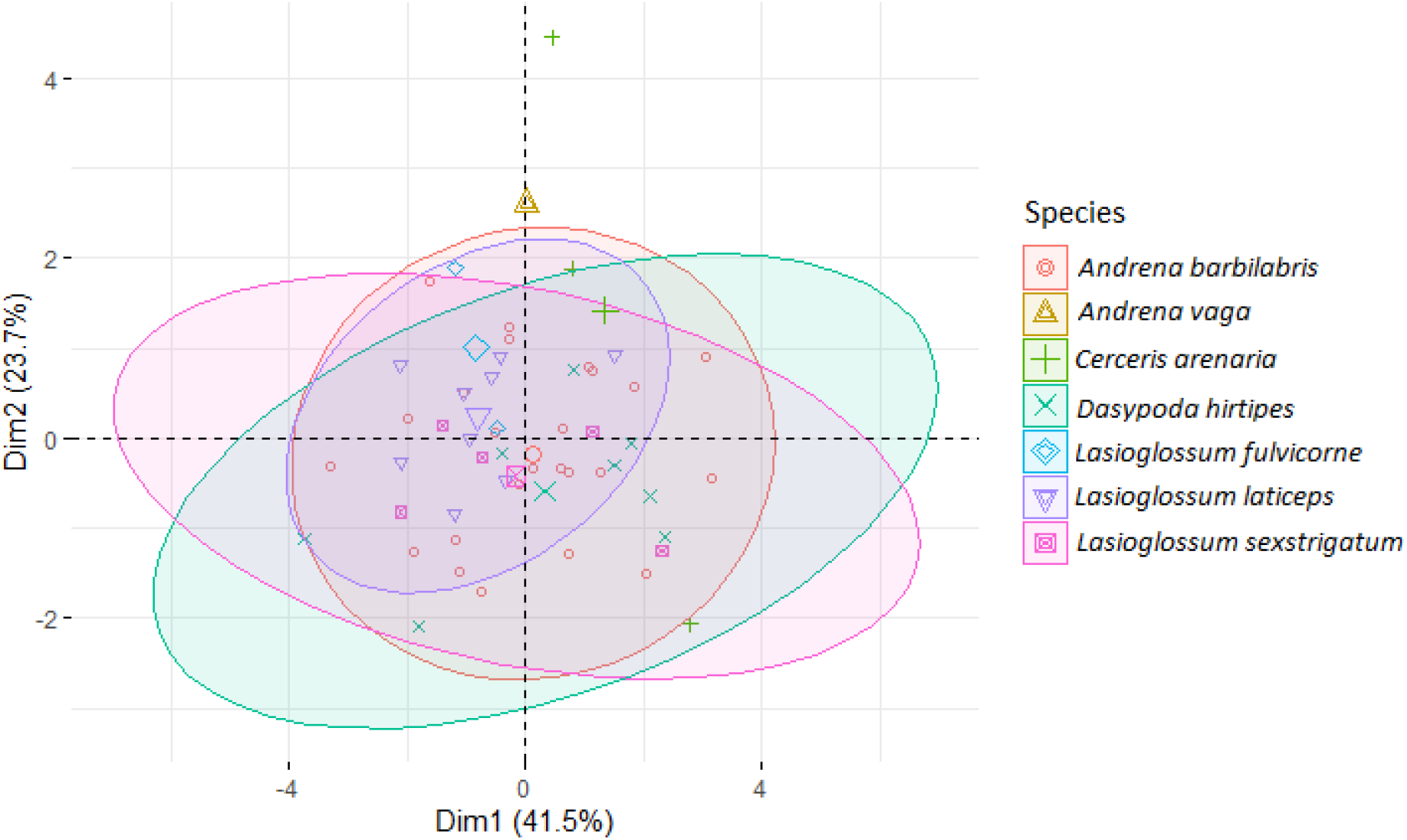
Principal component analysis (PCA) of collected tumuli samples grouping with 80% confidence ellipses by ground-nesting species. Dimensions 1 and 2 showed 65.2% of the explained variance. Coloured and shaped points (N = 53) corresponded to the ground-nesting species.

## Discussion

### Participatory survey and nest location

As data survey were heterogeneous on spatiotemporal aspect, the use of non-protocoled data from citizen science generated variations in the spatiotemporal prospection strength (Royle et al. 2005; Giraud et al. 2016). The disparity in recording frequencies could be partly explained by the implementation of the lockdown in mid-March 2020 related to the covid-19 health crisis that strongly restrained BRC citizens from leaving their homes and then, from locating and reporting potential nesting sites. Conversely, the increase in BCR temperatures and the rate of dissemination of the form over time, as well as the gradual unlocking of the BRC citizens in the weeks that followed, likely had a positive impact on the rate of form responses. The spatiotemporal nest observations could also be the result of divergences between the communication campaigns of the municipality administrations and the receptivity combined with socio-economic profile of the BRC citizens. These questions must be studied in greater depth by means of a regional social geography study.

Finally, we wondered how the representativeness of the revealed sites was in relation to the real number of nest-aggregations that cover the pavement of BRC and then on the real variability of sites captured by our study. Indeed, the heterogeneity of site detectability represented a common bias in participatory science (Royle et al. 2005; Kéry et al. 2009; Dorazio 2014). Nests located in crowded streets, housing large nest-aggregations as well as large sized identified species such as *Andrena* spp., *D. hirtipes, P. triangulum* or even *Cerceris* spp. (Table 1) with their respective sandy mounds were more prone to be detected. While small species nesting within the pavements such as *Lasioglossum* spp. or the other small-sized apoid species (Table 1) will be easily undetected because they were located in little frequented streets, sheltering small nest-aggregations with invisible sandy mounds. We can therefore assume that large species could be overestimated while small species could be underestimated in our sampling effort. Multi-counts, also common in participatory science processes (Guélat and Kéry, 2018; Renner et al., 2015; Warton et al., 2013), were discarded via the comparison of input lines.

### Monitored species

In our study, we were able to confirm some observations by Pauly in 2019a for *D. hirtipes, L. laticeps* and *A. barbilabris* (the latter also having been mentioned by Falk (2015)) but not for the Andrenidae bee species *Panurgus calcuratus* (Scopoli 1763). We may have missed the presence of this oligolectic specimen of *Heriacium* spp. (Asteraceae) because of its summer phenology. Its activity period corresponds to the end of June to the end of August (Rasmont and Haubruge 2002). However, some species of ground-nesting bees were identified for the first time nesting under the urban pavements: *A. vaga, L. sextrigatum* and *L. fulvicorne*. This total of 6 species of ground-nesting bees are among the common species and not concerned by an extinction status in Belgium with stable populations except for *A. barbilabris* whose populations are reported to be increasing. Moreover, there are no major conservation issues for these species in Europe (Nieto et al. 2014; Drossart et al. 2019). They are all polylectic except for *A. vaga* and *D. hirtipes* which are specialist bees of *Salix* spp. (Salicaceae) and some genera of Asteraceae respectively (Rasmont and Haubruge 2002). The 5 species of cuckoo bees such as *N. alboguttata* and *Sphecodes* spp. do not also have an extinction status in Belgium (Drossart et al. 2019). Compared to co-occurrence data with their associated hosts (Fig. S3), it would appear that *L. laticeps* could be a new host for the cuckoo *S. crassus* although we cannot confirm this by an entry in the nest. We also captured a specimen of *S. monilicornis* which would be a potential cuckoo bee of *L. laticeps* (Bogusch 2003). Moreover, we can support Vegter (1993) observations of large numbers of *S. miniatus* parasiting *L. sexstrigatum* nests. Lastly, three specimens of *S. pellucidus* and five of *N. alboguttata* were captured at sites of its known host *A. barbilabris* (Witt 1992; Rasmont and Haubruge 2002).

To our knowledge, ground-nesting wasps have never been recorded in urban pavements in the scientific literature. Our document is therefore the first written mention of these species nesting under urban pavements. Surprisingly, the species richness of these apoid wasps, all belonging to the Apoidea super-family, is greater than the identified ground-nesting bees. The sampled apoid wasp species have a solitary behaviour even though adults can also nest with a substantial density of conspecific (Bitsch and Leclercq 1993). Apoid adult wasps are generally nectarivorous but also capture preys by paralyzing them for their offspring development.

*P. triangulum*, commonly known as the “beewolf”, is a predatory apoid species that is widespread in Europe. It is a specialist predator of *Apis mellifera* L. but can exceptionally substitute on other genera of wild bees (e.g. *Andrena* spp., *Dasypoda* spp.) (Bitsch et al. 1997). The three *Cerceris* species are also solitary wasps nesting in sandy substrates. For their offspring, they captured and paralyzed small Coleoptera, notably Curculionidae or Chrysomelidae specimens for *C. arenaria* and *C. quadricincta* while *C. rybyiensis* is specialized in the capture of small species of halictid bees (Bitsch et al. 1997). *H. gerstaekeri* and *H. nobile* are known as nest parasitoid wasps. At present, there is no red list of Apoidae wasp species in Belgium, but we noted that *C. quadricincta* was included in the red list of endangered species in Germany (Haeseler and Schmidt 1984).

Surprisingly, other species of apoid wasps belonging to other genera, smaller and captured in lesser abundance, were also among the pavement’s inhabitants. These were *G. planifrons, D. insidiosus, M. lutaria, O. bipunctatus* and *L. pygmaeus armatus*. Their nests often consisted of a single gallery leading to the larval cell(s) in sandy soils. As *Cerceris* spp., the adults are generalist predators. Their preys consist mainly of specimens belonging to the families Cicadellidae, Fulgoridae, Cercopidae, Membracidae, micro-Diptera or micro-Hymenoptera (Bitsch and Leclercq 1993; Bitsch et al. 1997, 2007). It corresponds to common species in Belgium (Bitsch et al. 1997) and have no conservation status in Belgium even if *G. planifrons* is red-listed in Germany (Haeseler and Schmidt 1984). However, *L. pygmaeus armatus* is an uncommon psammophilous species in Belgium, with only two observations since 1950 despite its wide European range (Bitsch and Leclercq 1993; Rasmont and Haubruge 2002). This species was observed on 4 different sites in BRC, which would indicate that it nests commonly in BRC pavements and could therefore be the subject of a conservation project. Another finding of our study was that the investigated nest aggregation is species-mixed such as *C. arenaria* and *D. hirtipes* in a large series of sites (Fig. S3). From our sample protocol, it was difficult to observe nesting distinctions or sharing between ground-nesting species. Moreover, intraspecific specimens of *Cerceris* species can co-occupy the same nest (Willmer 1985; Polidori et al. 2006), which questioned about their strictly solitary behaviour. Therefore, it would be interesting to excavate the nest and observe the sharing of their larval cell structure.

Moreover, our observations have shown that urban pavements can also host another cohort of taxa that cohabit within the same segment of sidewalk, such as colonies of ant species. Ant colonies were not identified in this study although we observed that their nests seem to be more abundant in pavement compared to ground-nesting wasp and bee populations. Some of these ant colonies belonged to *Tetramorium* spp. (L.) (Stéphane DeGreef, personal communication). Since ants act as ecosystem engineers providing a great variety of ecosystem services (Folgarait 1998; Del Toro et al. 2012), a study seems necessary to shed light on the potential diversity of Formicidae nesting in BRC sidewalks.

Our specimen collection only reflected the captures over a 30 - 45 minutes period. It was likely that the whole diversity per site was not sampled. It should be noted that nest-aggregations were dynamic systems and the counts taken only reflect a moment in time in the life of ground-nesting insect populations and communities, suggesting that we probably missed species and nests that emerged at other period of the year. From a spatial point of view, this study was only framed in the Belgian capital under its own urbanization and climatic conditions. Therefore, extending this study to other large cities (e.g., Paris, Berlin…) could highlight other species of ground-nesting bees and wasps in urban pavements and which potentially have different conservation issues.

### Joint size and pavement structure

Given the newness of the study of pavement nesting sites of ground-nesting species, no scientific references were available to compare our results with current or previous studies on this topic. For this reason, many hypotheses and questions were raised in this discussion part. The joint size usually designed in Brussels vary from 1mm to 1.5cm depending on the shape of the pavement element and the maximum diameter of the jointing material (Bruxelles-Mobilité 2016). However, during our observations, we determined that the size of joint hosting gallery entrances oscillated around a wider average of 1.08cm with a standard deviation of 0.57cm without any real distinction between the species or families of Apoid. An increase in size between two tiles or pavers is likely to lead to pavement degradation. This suggested that these species of ground-nesting bees and wasps tended to nest in pavements with relatively wide joints by the standard specifications used in BCR (Bruxelles-Mobilité 2016). Furthermore, our results showed that joint size was not a function of individual size, implying that ground-nesting species did not have a preferential size according to their size. Probably, the morphological limit for Apoid to pass through a given diameter is the distance from the tips of their thorax. The thoracic cuticle should be thicker and harder than the abdominal cuticle (Elias-Neto et al. 2009). Among all the sampled species, ITD varied between 0.20cm and 0.45cm which was sufficient to allow them to pass through the narrowest measured pointing.

All sites showed an opening at the joint level which allowed the ground nesting species to dig their galleries. The observed joint openings were either the result of the absence of jointing material, or of the presence of unbound jointing material or of the degradation of a bound jointing material. The presence of galleries at the start of bound joints highlighted in the results was rather surprising. Indeed, these structures were in principle completely closed and did not allow insect nesting. We observed simultaneously pavement degradation and jointing material fragmentation. This could be explained by the poor quality of the rigid joint which impacted the durability and cohesion of the material and made it more prone to disintegration during any disturbance (shrinkage cracking, freeze/thaw episodes, etc.). Also, the age of the jointing material could also have an influence on its current state of deterioration. Another potential scenario would be whether these ground-nesting species were involved in the degradation of the modular structures and in particular their joints, or whether they took advantage of the presence of a previously degraded structure to nest, or a combination of the two hypotheses. Bees are able to dig into hard-packed soils (Barthell et al. 1988; Cane 1991). Moreover, a BCR pavement with unbound jointing materials is theoretically always coupled with underlying permeable and draining layers (i.e., sand, gravel, stonework) in order to avoid water stagnation in the structure as well as its deterioration (CRR 2009, 2018). This combination of materials is known as unbound flexible paving and concerns nearly 80% of the sites encountered in our study. It seems to be in line with the criteria of sandy texture and drainage of the soil material generally required by ground-nesting bees and wasps in their natural environment. Indeed, as Wuellner (1999) pointed out, a soil that was too waterlogged, flooded or too dry can jeopardise the survival capacity of individuals in the larval and immature stages. Moreover, the presence of ground-nesting bees on the same segment of pavement for several years suggests reproductive success and optimal conditions for perennial occupation within urban pavements. In the context of this work, only the texture of the sand present at the interface between the sub-base and the foundation was studied (see supplementary information 1) and it could be interesting to also study the composition of the substrate both at the level of the joint and the sub-base and the other strata which make up the pavement.

Most pavements were composed of sandstone setts and concrete slabs. This suggested that pavement thickness and type had little effect on nesting site preference for any ground-nesting species. These findings raised many questions about a possible multiplicity of gallery forms within the substrate. Even though we were able to determine the depth at which we found individuals of *A. barbilabris* at 20 and 27 cm (see supplementary information 1), did the tunnels plunge deep into the different layers that make up the pavement or do they extend to the first few centimetres under the pavement ? In addition, we can wonder whether the choice of nesting location resulted from a preference for concrete slabs and sandstone setts in particular, or whether their location was simply the result of a higher availability of these types of paving in BCR. Our study did not investigate these questions in depth, but they would nevertheless merit further investigations.

As for the results relating to the location of the sites on the pavement, they enabled us to highlight phenomena that had not been recorded to our knowledge in the literature until now. Indeed, although our observations suggested that many individuals nested on pavements only, we also observed nesting sites at the level of house steps and stairways. This observation was accompanied by a lack of jointing material along these terraced houses, which allowed easier access to the sandy stratum under the pavements for ground-nesting species. It was mainly *Lasioglossum* spp. and non-Philantidae species that preferred to nest in this type of location, which also should allow them to benefit from the heating of their nest entrance by solar radiation (Cane 2015) that first reached the facades of the terraced houses. This absence of jointing material along houses was common in the city, which suggested that the real abundance of these species, especially *Lasioglossum* spp., had probably been underestimated in BCR. According to our field observations, these halictids were therefore the main inhabitants of the sandy stratum under the pavements.

All the observations in the joint size and pavement structure highlight that the mineralization and coating of surfaces in urban areas was not always correlated with a net loss of nesting opportunities as suggested by Cane et al., (2006) and Fortel et al., (2016). On the contrary, our study highlighted the opportunity that these structures can represent in terms of nesting resources in the city. Finally, our observations tend to confirm the hypothesis put forward by Pauly (2019a), who stated that among all pavement types in BRC, old pavements in BCR - where the soil under the paving stones was sandy and where the joints were not cemented - were the most hospitable for ground-nesting bees and consequently for ground-nesting wasps as well. Nevertheless, the old BCR pavements were more prone to be redeveloped by project owners which should destruct existing nests.

### Soil texture analysis

The particle size analysis revealed that the mound samples corresponded to the sandy texture in the USDA triangle. This texture class was consistent with observations made by Cane (1991) on 32 species of ground-nesting bees in the USA, by Vereecken *et al*., (2006) for *A. vaga*, Malyshev (1935) and Michez (2007) for *D. hirtipes*, and Falk (2015) on the 6 remaining bee species, which stipulated that these ground-nesting bees built their nest in particular on sandy-textured soil. However, while these authors noted a multitude of other used textures, such as silt loams and clay loams for Cane (1991), sandy-clay soils also for Vereecken *et al*., (2006) or clay soils also for Michez (2007), our study highlighted only one type of texture used by the bees to dig their galleries. This means that the foundation layer could be homogeneous across all BRC pavements if it had a sandy origin. On the other hand, encouraging a sandy and homogeneous texture through urban redevelopment would only filter specific populations related to this ecological niche.

In addition, certain parameters such as vegetation cover, percentage of organic matter, drainage, temperature, exposure and soil aspect were not studied in this work but have been identified in the literature as factors impacting on nest selection (Stephen 1960, 1965; Osgood Jr 1972; Potts and Willmer 1997; Wuellner 1999; Sardiñas and Kremen 2014; Cane 2015; Anderson and Harmon-threatt 2016; Harmon-Threatt 2020; Nichols et al. 2020). It would therefore be interesting to study these parameters in the future, to assess global trends and to compare these characteristics by species in order to increase knowledge on this subject, which is currently poorly known (Cope et al. 2019; Buchholz and Egerer 2020; Antoine and Forrest 2020). Even fewer studies have been carried out on the edaphic preferences of apoid wasps, even though our results showed that these species followed the soil texture preferences of ground-nesting bees.

### Implications for urban pavement design and their management

Suitable pavement for ground-nesting species consisted of sandstone pavers or concrete slabs with an unbound jointing size around 1cm on an unbound foundation. However, this pavement hosting ground-nesting species was only an archetype (Fig. 7) from the results of our study to BCR specifications. We considered other alternatives for urban pavements that would probably host urban insect community. For potential conservation, it is important to provide a structure that is favourable for new nesting process. In general, common practices recommend that joint size should be as small as possible, which should limit the access for the ground-nesting species. It therefore seems difficult to modify the requirements of the standard specifications established by BCR specifications, as these are the basic rules for guaranteeing the durability of the pavement (Bruxelles-Mobilité 2016).

**Fig. 7.**
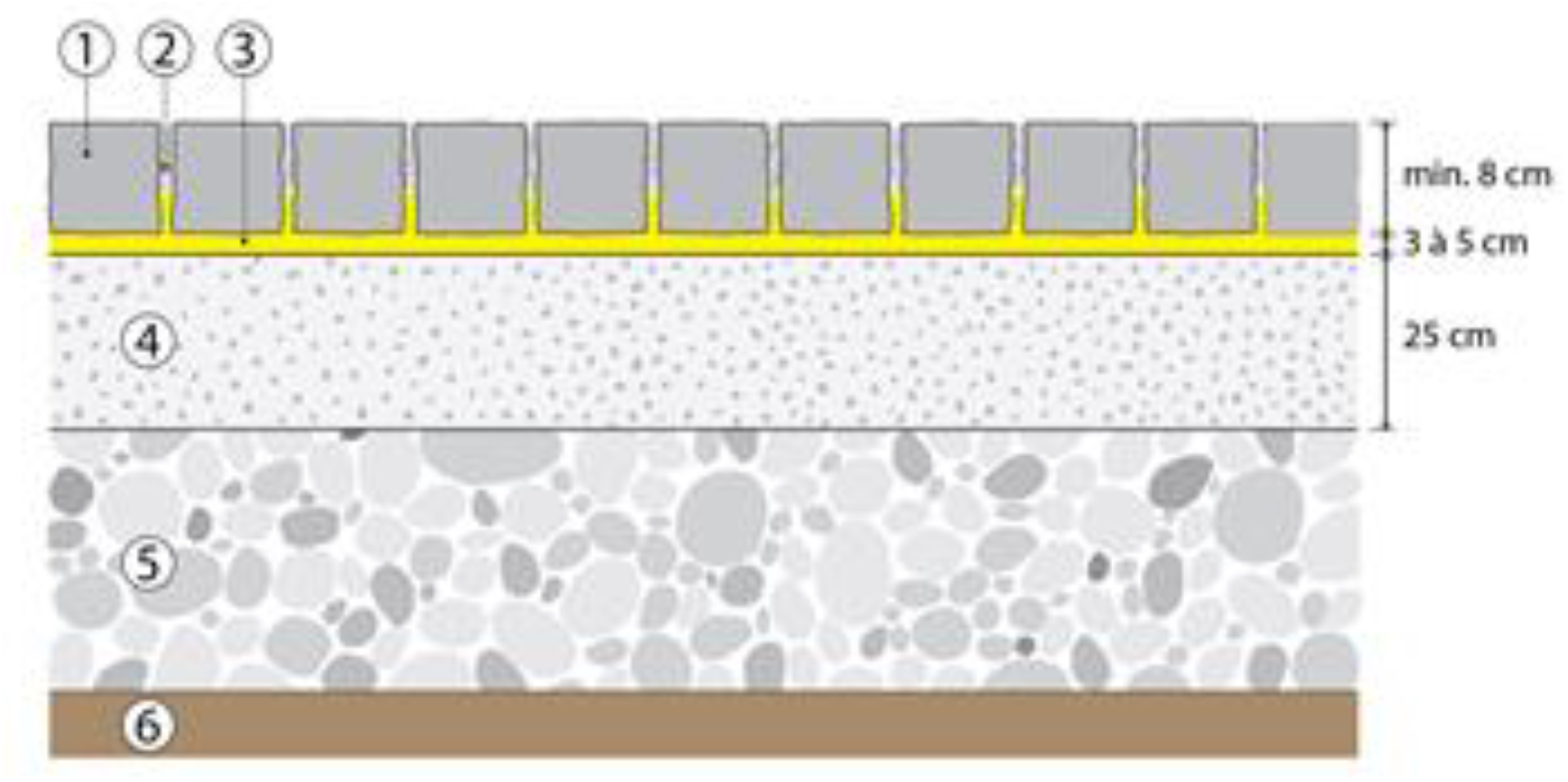
The pavement structure : (**1**) Paving elements ; (**2**) Joints with an opening size of 0.8 to 1.0cm, filled with sand 0/6.3 or 0/8 (fine content less or equal than 10%); (**3**) Laying course: gravel 2/6,3 or 2/8 (fine content less or equal than 2%); (**4**) Road base: unbound aggregate 0/20 or 0/40; (**5**) Sub-base with defined thickness according to the construction plan; (**6**) Subgrade

Moreover, natural stone pavements with wide and unbound jointing material were therefore more favourable to the nesting of the studied apoid species even if they generally offer a low level of pedestrian comfort (Bertrand et al. 2019) and are not totally compatible with the durability requirements from BCR specifications which recommend modular structures of the bonded type and with joint opening smaller or equal than 1cm (Bruxelles-Mobilité 2016). This finding highlights a discrepancy between the societal requirements of BCR and the ecological opportunities.

Therefore, we suggest new pilot studies to be carried out to assess the possibility of designing multifunctional pavements that could simultaneously meet the challenges of comfort, durability, hosting entomo-biodiversity and rainwater infiltration. Either by means of technological advances on paving bloc design, or by differentiating pavements with a central zone dedicated to pedestrians (no joints or thin joints) and aside zone dedicated to the reception of ground-nesting species and the infiltration of water (wide joint openings, draining material, vegetated or not), or by creating and/or maintaining a vegetated strip peripheral to the pavements as they can exist in BCR allotments (Fig. S6).

In addition, we recommend the implementation of localized campaigns to raise awareness of urban ground-nesting bees and wasps among resident citizens in order to avoid nest destruction due to a lack of knowledge of the ecology of these species as they are solitary and not dangerous to humans. Similarly, it would be interesting to ensure continuous actions in order to initiate and reinforce a change of paradigm with regard to the city’s aesthetic criteria: “clean” and “unfunctional” pavements in human-centered city vs. “multifunctional” pavements in bio-centered city (Aronson et al. 2017; Rivkin et al. 2019).

## Conclusions

This study confirmed the location of 89 sites of ground-nesting bee and wasp species nesting under pavements spread over almost BCR. Surprisingly, a higher species richness than expected was identified for a total of 22 species nesting in these pavements, whose one Crabronidae species, *L. pygmaeus armatus*, could benefit from conservation issues. Several types of nest aggregation were also identified. A part of them hosted a single species of bee or wasp, while others were composite. The sandy textural layer under the pavement as well as its access -conditioned by the nature of the joints-seem to be the main factors influencing the selection of nesting sites for these ground-nesting species. BCR pavement thickness and type have little effect on nest site preference for all species. Old BCR pavements made of natural stone pavers or concrete slabs with wide, unbound jointing material are therefore more favourable for the sandy ground-nesting species. But, in terms of durability, comfort and stability, continuous or modular structures of the bounded type and with narrow joints are currently the most recommended by BCR specifications and therefore highlight an overhang between the BCR societal requirements and the ecological opportunities of the sampled species.

This study pointed out that pavements, previously perceived as inappropriate for biodiversity, could serve as refuge for some insect populations. Therefore, by modulating pavement design and proposing alternative construction models, we could turn the city into a more welcoming place for biodiversity.

## Supporting information

Supplementary Informations

## Statements & Declarations

### Author Contributions

All authors contributed to the study conception and study design. Material preparation and data collection were performed by Grégoire Noël, Violette Van Keymeulen, Sylvie Smets and Olivier Van Damme. Analysis was performed by Grégoire Noël and Violette Van Keymeulen. The first draft of the manuscript was written by Grégoire Noël and all authors commented on previous versions of the manuscript. All authors read and approved the final manuscript.

## Acknowledgements

Thanks to Julie Bonnet (ULiège, Functional and Evolutionary Entomology) for the insect preparation and some identifications. The R code and the data used in this paper are available at https://github.com/gregnoel/Street_Apoid.git

