## Supplementary Informations for "Nest aggregations of wild bees and apoid wasps in urban pavements: a “street life” to be promoted in urban planning"

**Supplementary information 1:**

**Supplementary soil texture method**

We dismantled several modular elements from pavement structure, assessed the nature of the laying course and, where possible, excavated its foundation to determine its nature and thickness. A 250-300g sample of sand from two sites was taken for grain size determination.

The two sand samples taken on site were sieved in BRRC's laboratories according to the requirements of NBN EN 933-1 and 2, applied in road engineering. A dry sieving was carried out. The opening of the sieves was chosen to be as close as possible to the sieves applied in pedology, which are slightly different from those applied in road engineering. Thus the 100µm sieve was replaced by the 90 µm sieve and the 50 µm sieve by the 63 µm sieve.

**Results of the excavation**

1) The description of the first excavation (concrete slabs in Watermael-Boitsfort, Bruxelles):

The layer course consisted of a thick layer of mortar, contrary to what was expected in the presence of sandy joints (sometimes mixed with mortar). The removal of the pavement was limited to one slab in order not to destabilize the structure. As a result, it was not possible to excavate the foundation to determine its thickness. The concrete slabs (30 x 30 cm) probably laid in a full mortar bath, show joints filled with mortar (or damaged) or sand.

The joint width was very irregular, between 1 and 15 mm (average 8mm) and the thickness of the mortar layer was irregular at 3-4 cm. We observed a layer of yellow sand (foundation or laying course).

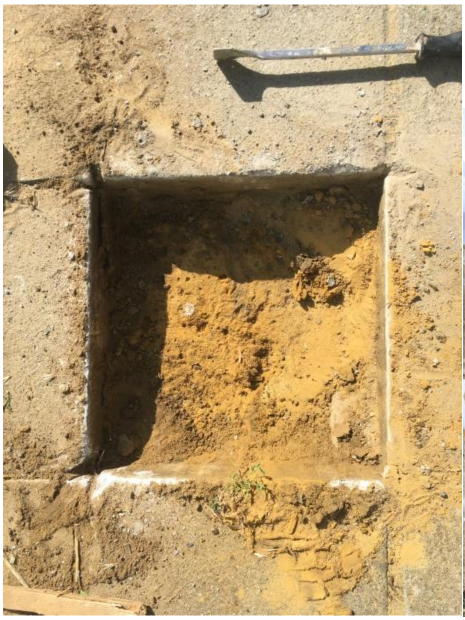

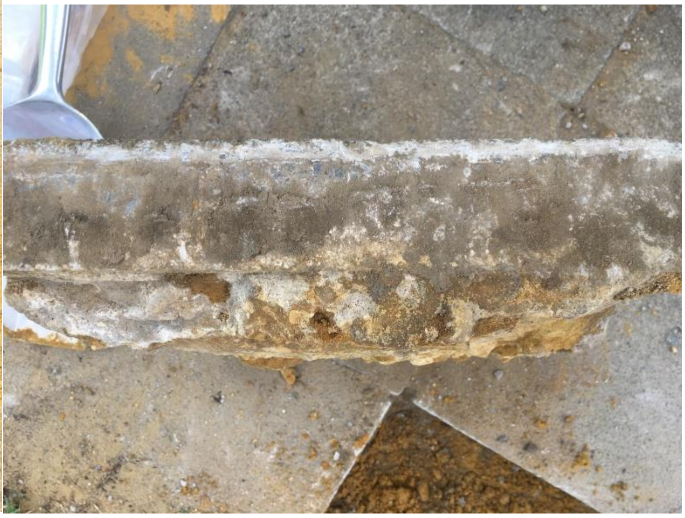

*First excavation (Watermael-Boitsfort, Bruxelles) - Left: Removal of the concrete slab. Right: Concrete slab and profile mortar thickness. Images by Sylvie Smets*

2) The description of the second excavation (sandstone setts in Etterbeek, Bruxelles):

The sandstone setts (12cm * 12cm) have a thickness of 9 cm and are jointed with sand. The width of the jointing was on average 5mm. The laying course with a thickness of few centimeters is composed of yellow sand. The limit between the laying course and the basis was unclear. The foundation is composed of aggregates with some brick debris, with 35 cm thick, including the laying course. The sub-base or soil was composed of indurated material and could not be identified.

We also observed the presence of *Andrena barbilabris* (2 living individuals) found at 20 and 27 cm depth, the presence of *Lasioglossum laticeps* (1 dead individual) found at 7 cm depth, the presence of ants (5 living individuals) found at different depths. We also discovered some chambers containing weevil corpses at depth of 10, 20 and 30 cm with eggs leading to the hypothesis of the presence of *Cerceris* spp. (probably *Cerceris arenaria*).

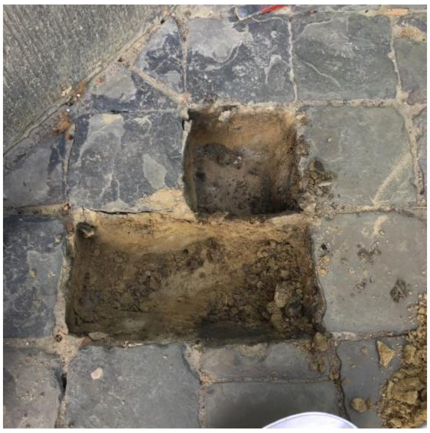
1

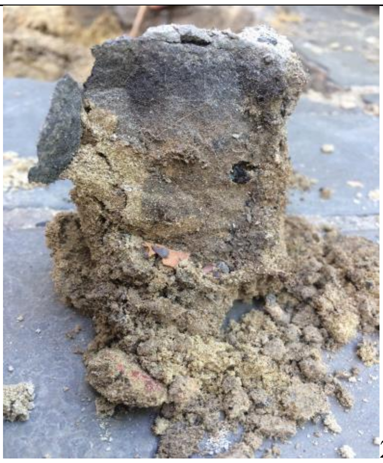
2

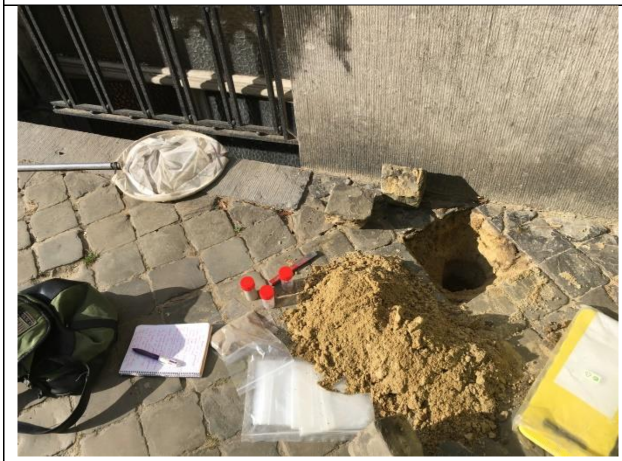
3

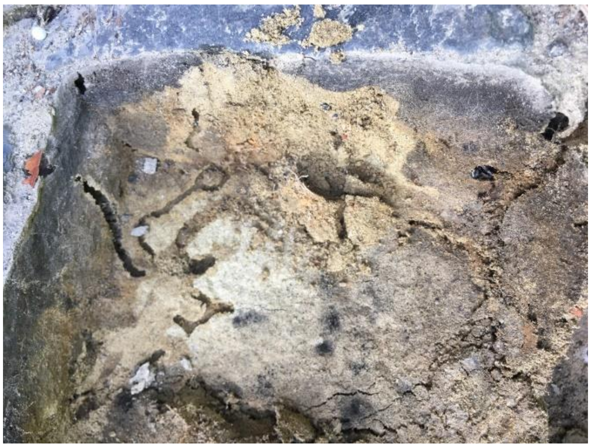
4

| | |
|---|---|
| 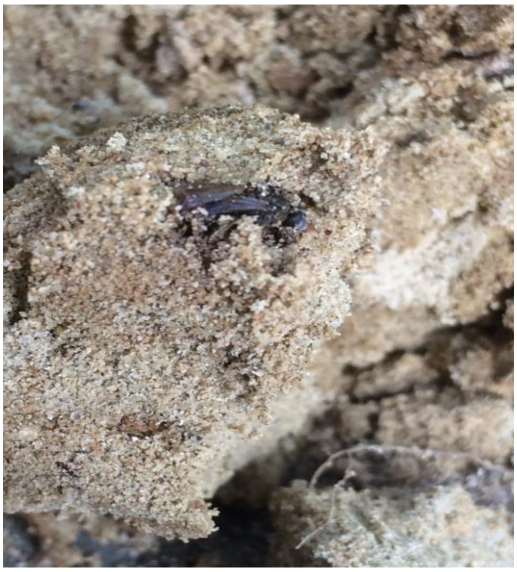 5 | 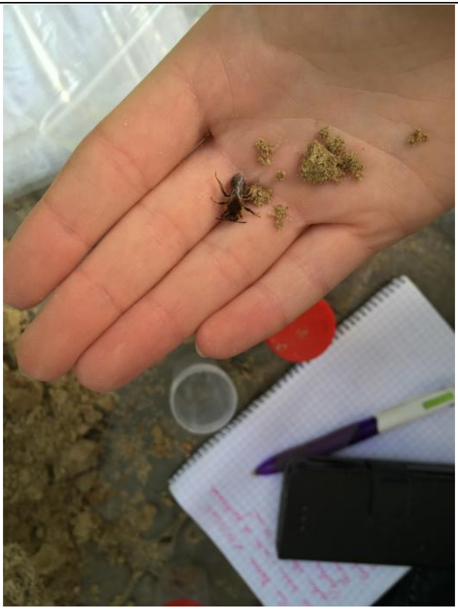 6 |

*Second excavation (Etterbeek, Brussels) - 1: Removal of paving stones; 2: Profile of the first few cm; 3: Excavation of the yellow sand basis; 4: Gallery in the basis layer; 5: Weevil corpse; 6: Andrena barbilabris in the basis (20-27 cm deep). Images by Sylvie Smets (CRR) and Violette Van Keymeulen (ULiège).*

**Measurement of the grain size of the samples taken on site**

The particle size analysis of the two sand samples taken on these sites could be comparable to those carried out on the mound sand, given the slight difference in the limits of the classes (90µm instead of 100µm and 63 µm instead of 50 µm). We noted that the most represented category of base layer sands is the 200 to 90 µm class, whereas for the mound sands, it is the 500 to 200 µm class that is most represented. It should also be noted that the measurements on mound sand are much more numerous. We also noted that the percentage of material below 63µm is 4.5% on average. There is therefore less than 4.5% of the fraction (silt + clay, limits 2 and 50µm) and we are therefore in a predominantly sandy texture, as for the materials recovered from the sandy mounds at the surface of the pavements.

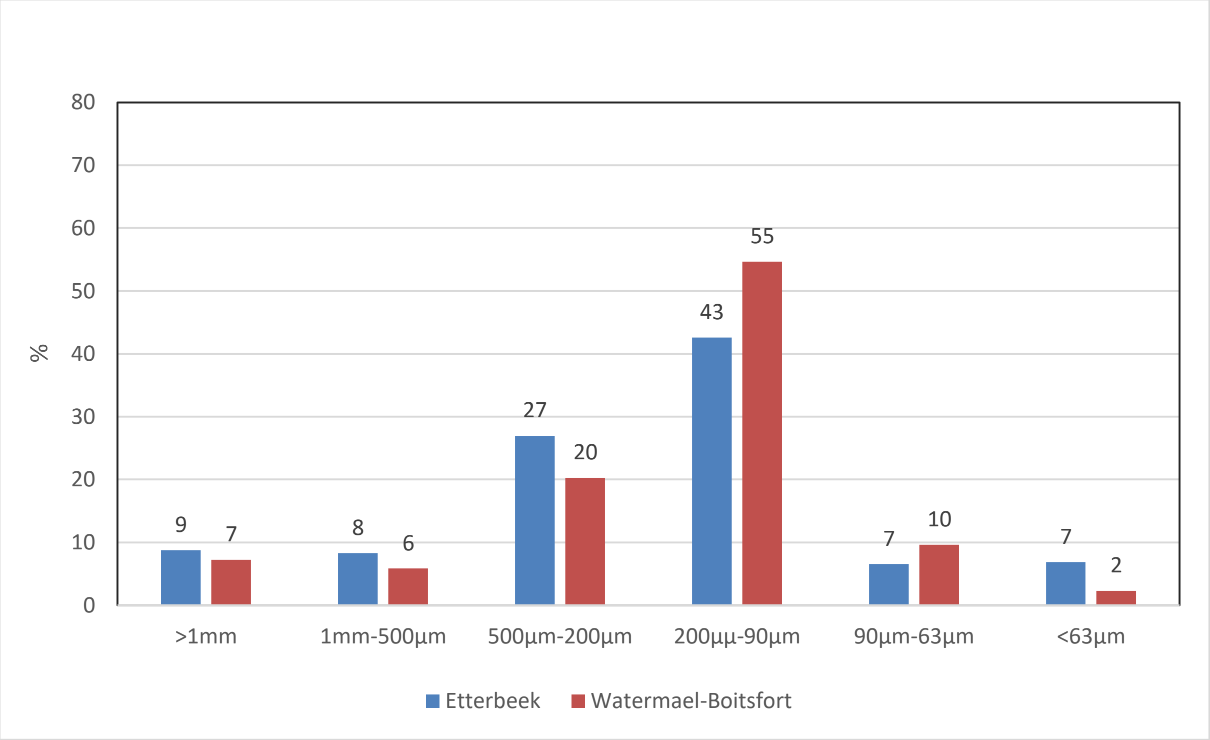


*Size distribution of sand taken from under the mortar layer (Watermael-Boitsfort) and laying course (Etterbeek) taken on site.*

**Participatory survey**

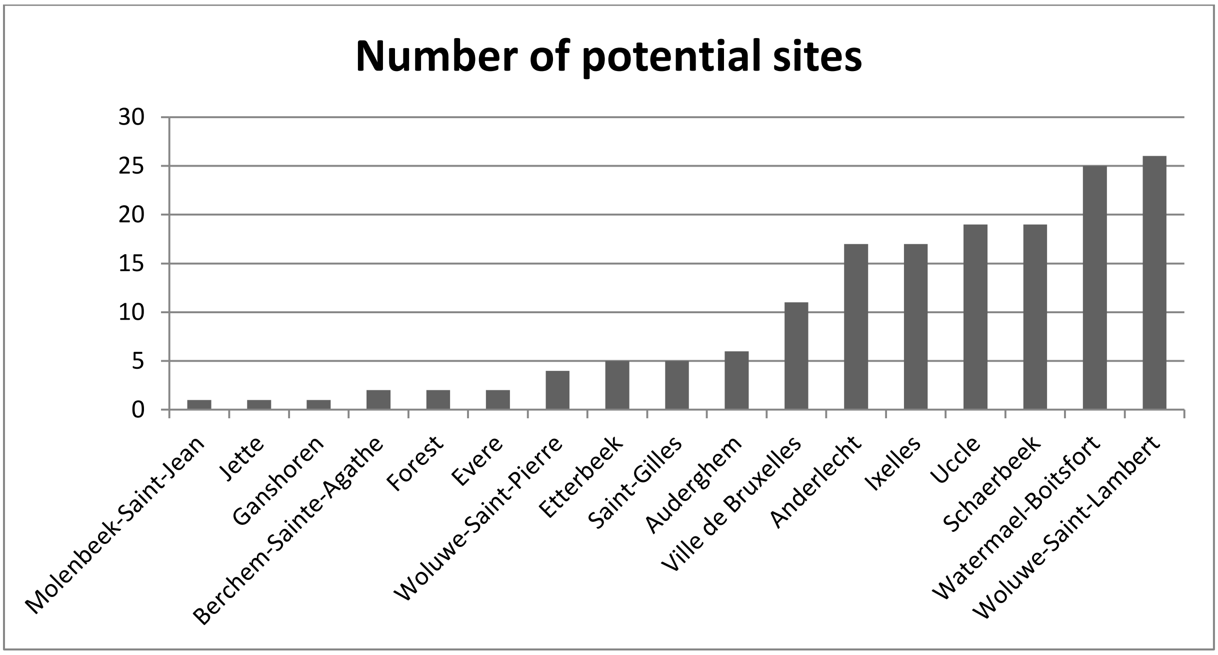


*Figure S1 – Spatial distribution of potential sites by municipality using the coding form*

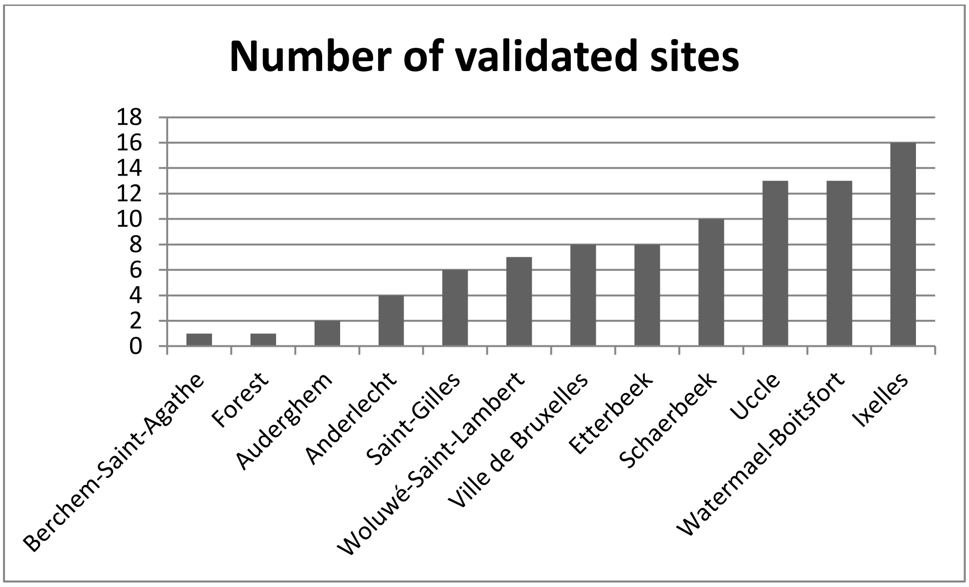


*Figure S2 – Spatial distribution of validated sampling sites by municipality*

**Species recorded**

We observe that all species are linked by at least one link to each other except for *A. vaga*. The most pronounced co-occurrences for ground-nesting species are between *C. arenaria* and *L. pygmaeus*, between *C. rybiensis* and *L. laticeps*, between *A. barbilabris* and *L. laticeps*, and between *D. hirtipes* and *C. arenaria*. The co-occurrence between cuckoo species and their associated hosts are also present: between *H. nobile* and *C. arenaria*, between *H. gerstaeckeri* and *M. lutaria*, between *H. gerstaeckeri* and *C. arenaria*, and between *S. crassus* and *L. laticeps*.

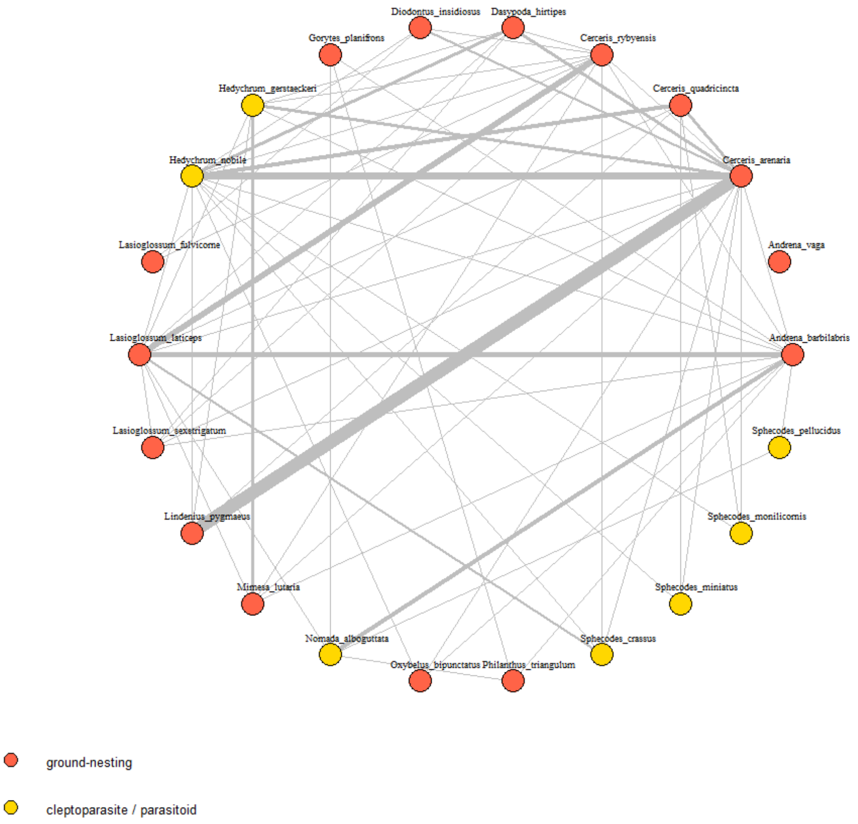


*Figure S3 – Co-occurrence network for identified bee and wasp species. The red and yellow colours represent their nesting strategy: ground-nesting or cleptoparasitic/parasitoid. The link (grey line) between two species means that they have shared the same site at least once. The size of the linkage is dependent on the number of times these species cohabit on the same sites.*

The number of nests present on the sidewalk shows great variability and ranges from 2 to 500 nests with an average of 107 nests per site and a median of 50 nests per site. This range between the median and the mean indicates an asymmetric distribution with a small number of data with high values. As for the nest density, it presents a mean of 12 nests/m² and oscillates between 0.167 and 100 nests/m². It shows an asymmetric distribution rather similar to the number of nests with a median at 5 nests/m².

**Joint Size Analysis**

*Table S1 - Measurements of the joint sizes (width) near the nest entrance of ground-nesting species in urban pavements.*

| Species | Family | Measure number | Mean size ± standard deviation [cm] | Size min/max [cm] |
|---|---|---|---|---|
| *Andrena barbilabris* | Andrenidae | 145 | 1,20 ± 0,62 | 0,3/3,0 |
| *Andrena vaga* | Andrenidae | 3 | 0,7 ± 0,1 | 0,6/0,8 |
| *Cerceris arenaria* | Crabronidae | 110 | 1,04 ± 0,51 | 0,4/3,0 |
| *Cerceris quadricincta* | Crabronidae | 9 | 0,91 ± 0,23 | 0,6/1,3 |
| *Cerceris rybyensis* | Crabronidae | 7 | 1 ± 0,56 | 0,5/2,2 |
| *Dasypoda hirtipes* | Melittidae | 29 | 1 ± 0,34 | 0,4/3,0 |
| *Lasioglossum fulvicorne* | Halictidae | 11 | 0,82 ± 0,44 | 0,5/2,0 |
| *Lasioglossum laticeps* | Halictidae | 56 | 0,93 ± 0,63 | 0,2/3,0 |
| *Lasioglossum sexstrigatum* | Halictidae | 19 | 1,19 ± 0,75 | 0,3/2,5 |
| *Philanthus triangulum* | Crabronidae | 9 | 1,27 ± 0,51 | 0,8/2,5 |

### Granulometry analysis

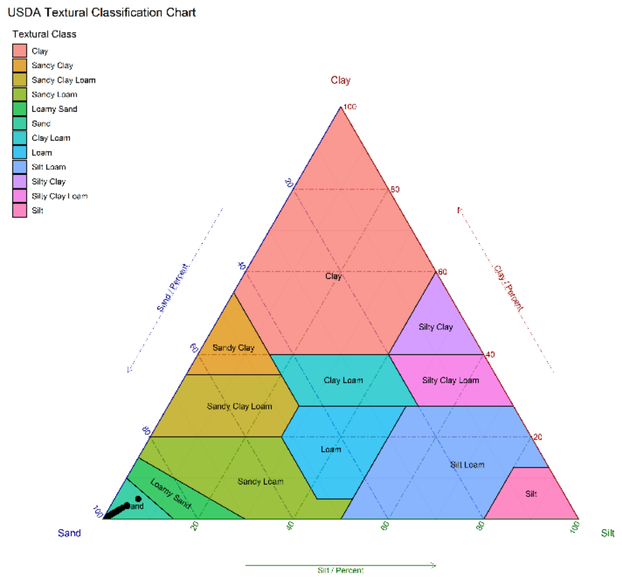


*Figure S4 – USDA textural triangle and the location of the collected tumuli samples in black dot points (N = 53)*

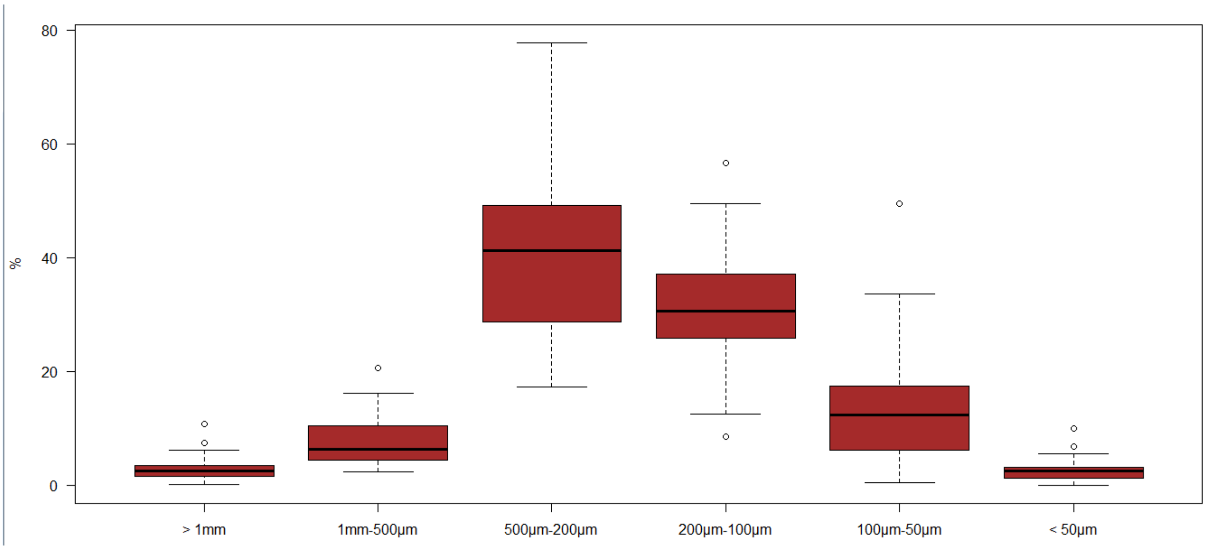


*Figure S5 – Boxplot of the quantitative variables related to the rates of particles size fractions of the sandy mounds of bees and wasps nesting in the pavements of the Brussels-Capital Region*

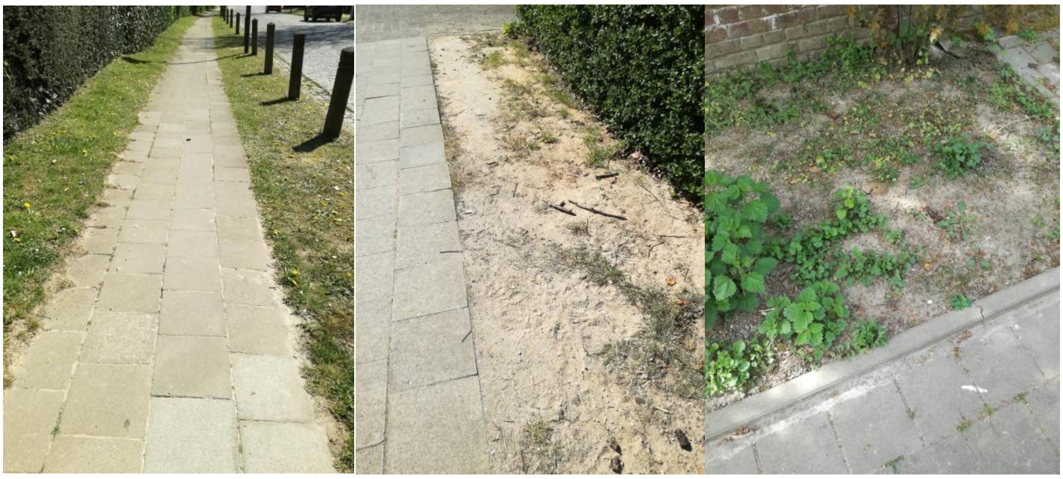

*Figure S6 – Pavements in the municipality of Uccle that have grassy or non-vegetated beds to accommodate species of ground-nesting Hymenoptera.*
